# Specific interactions with phospho-Ubls and allosteric conformational changes regulate the E3 ligase activity of Parkin

**DOI:** 10.1101/2023.11.21.568018

**Authors:** Dipti Ranjan Lenka, Shradha Chaurasiya, Atul Kumar

**Affiliations:** Department of Biological Sciences, Indian Institute of Science Education and Research (IISER) Bhopal, Bhopal, India, 462066

## Abstract

PINK1 and Parkin mutations lead to the early onset of Parkinson’s disease. PINK1-mediated phosphorylation of its substrates such as ubiquitin (Ub), ubiquitin-like protein (NEDD8), and ubiquitin-like (Ubl) domain of Parkin activate autoinhibited Parkin E3 ligase. The mechanism of various phospho-Ubls binding on Parkin and conformational changes leading to Parkin activation remain elusive. Herein, we determine the first crystal structure of human Parkin E3 ligase bound with phospho (p)-NEDD8, which shows that NEDD8 has evolved over Ub to bind and activate Parkin more robustly. X-ray crystal structures and supporting biophysical/biochemical data reveal specific recognition and underlying mechanisms of pUb/pNEDD8 and pUbl domain binding to the RING1 and RING0 domains, respectively. This new data also shows that pUb/pNEDD8 binding in the RING1 pocket causes allosteric conformational changes in Parkin’s catalytic domain (RING2), leading to Parkin activation. Furthermore, Parkinson’s disease mutation K211N in the RING0 domain of Parkin was believed to lose its activation due to loss of interaction with pUb. However, our data reveal allosteric conformational changes due to N211 that lock RING2 with RING0 to inhibit Parkin K211N activity without disrupting pNEDD8/pUb binding. This study would aid the design of small-molecule Parkin activators for the treatment of Parkinson’s disease.

## Introduction

The PARK2 and PARK6 genes encode the Parkin E3 ligase and PINK1 (PTEN-induced putative kinase 1), respectively. Parkin and PINK1 control the mitochondrial homeostasis process through autophagy (mitophagy) of the damaged mitochondria. Mutations in the PARK2 and PARK6 genes cause Parkinson’s disease (PD) ^1–7^.

Parkin is a member of the RBR family of E3 ligases and consists of an N-terminal ubiquitin-like (Ubl) domain followed by four Zn2+binding domains named RING0, RING1, inbetweenRING (IBR), and RING2 domains ^8–10^. Like other members of the RBR family of E3 ligases, the catalytic C431 (on RING2) is charged with ubiquitin in an E2-dependent manner before ubiquitin is transferred to lysines on its substrate ^11–13^. Parkin is an autoinhibited cytosolic protein ^8^. The E2 binding site on Parkin is blocked by the presence of Ubl on RING1. The short repressor (REP) element between IBR and RING2 also restricts E2 binding. RING0, which occludes the catalytic C431 on RING2, significantly inhibits Parkin activity ^8,10,14,15^.

The phosphorylation of S65 on Ubiquitin by PINK1 leads to the recruitment of Parkin to mitochondria, which is mediated by the enhanced interaction between RING1 of Parkin and phospho-ubiquitin (pUb) ^16–19^. The conserved S65 on the ubiquitin-like (Ubl) domain of Parkin is also phosphorylated by PINK1 ^20,21^, leading to the fully active Parkin conformation mediated by interactions between phospho-Ubl (pUbl) and RING0. The RING0 pocket (comprising K161, R163, and K211) and RING1 pocket (comprising R305, H302, and K151) bind to the phospho-S65 group of phosphorylated activators of Parkin ^22–27^. Recently, PINK1-mediated S65 phosphorylation of another ubiquitin-like protein, NEDD8, was shown to activate Parkin ^28^. While several crystal structures of Parkin with pUb are known ^22,23,26,27,29,30^, the structure of pNEDD8 and the Parkin complex is not available. Furthermore, it remains elusive whether and how the two look-alike pockets (on RING0 and RING1) of Parkin recognize different Parkin activators (pUbl, pUb, and pNEDD8) specifically or promiscuously.

Recent publications have suggested two pUb binding sites on Parkin, one in the RING1 pocket and the other in the RING0 pocket ^29,31^. The latter suggests a promiscuous nature of interactions between Parkin and phosphorylated ubiquitin-like proteins/domain. Furthermore, these publications demonstrated that the disease mutation K211N on Parkin results in loss of Parkin activation in the presence of pUb, leading to loss of Parkin localization to the mitochondria ^31^. Parkin construct lacking the RING2 domain was used to demonstrate the binding of pUb on both the RING0 and RING1 pockets, and K211N mutation led to the loss of pUb binding on the RING0 pocket ^29,31^. Also, a new feedforward pathway of Parkin activation was proposed wherein pUb binding is preferred over pUbl binding in the RING0 pocket ^29,31^. Interestingly, our recent work demonstrated that trans interactions between pUbl and RING0 are negatively affected by the RING2 domain of Parkin ^30^. The latter work used a method to capture pUbl and RING0 interactions in trans without requiring the removal of RING2 from the Parkin construct ^30^. The negative effect of RING2 on the binding of phosphorylated Ubl to the RING0 could help RING0 to specifically select between pUb/pNEDD8 and pUbl. Additionally, our recent work showed that phosphorylation of the Ubl domain and its interaction with the RING0 domain in trans play an important role in Parkin recruitment to impaired mitochondria ^30^. In light of recent publications, a detailed biophysical study is warranted to understand the mechanism of interaction between Parkin and its activators.

Herein, we solve the x-ray crystal structure of Parkin and pNEDD8 complex, which revealed the binding of pNEDD8 to the RING1 pocket where pUb binds. However, the structure shows new interactions between pNEDD8 and Parkin, which are mediated by favorable residues on NEDD8 that are not conserved on pUbl or pUb. The latter is further supported by quantitative ubiquitination assays, which show more robust activation of Parkin by pNEDD8 than by pUb. We also demonstrate that pNEDD8/pUb binds specifically to the RING1 pocket and does not cross-react with the RING0 pocket, unlike previously reported. We demonstrate that the specificity of the RING0 and RING1 pockets is mediated by the unique electrostatic surface charge distribution on Parkin and Ubl/Ub/NEDD8. We show that a mutation (N60D) on Ub that mimics the electrostatic charge distribution of the Ubl domain of Parkin changes the specificity of pUb (N60D) and allows it to bind with the RING0 domain. Our data also shows that, unlike the binding of pUbl in the RING0 pocket, which results in large conformational changes in RING2, the binding of pUb/pNEDD8 in the RING1 pocket leads to dynamic RING2 due to allosteric conformational changes. Furthermore, we demonstrate that disease mutation (K211N) on Parkin perturbs E3 ligase activity independent of interactions between Parkin and pUb/pNEDD8, unlike previously suggested. We also solve the structure of K211N Parkin and pNEDD8 complex, which reveals allosteric conformational changes in RING0 due to N211, leading to new interactions between RING0 and RING2. These new interactions restrict the movement of RING2 in K211N Parkin, leading to the loss of E3 ligase activity independent of pUb/pNEDD8 binding. Overall, this study would aid therapeutic efforts to develop small-molecule activators against Parkin.

## Results

### NEDD8 has evolved over ubiquitin to bind and activate Parkin

Recent publications have suggested two pUb binding sites on Parkin, one in the RING1 pocket and another in the RING0 pocket where pUbl binds ^29,31^. Another study showed that NEDD8 phosphorylation by PINK1 leads to Parkin activation ^28^. While the phosphorylation of different ubiquitin-like proteins by PINK1 and their binding to Parkin are well known, the selectivity and specificity of the above pathway remain elusive. To test the specificity of the interaction between Parkin and pUb or pNEDD8, we performed quantitative ubiquitination assays on Parkin using increasing concentrations of pUb and pNEDD8. Interestingly, we noticed that the degree of pNEDD8-mediated Parkin activation was significantly greater than that of pUb-mediated Parkin activation (Fig. 1A).

**Figure 1.**
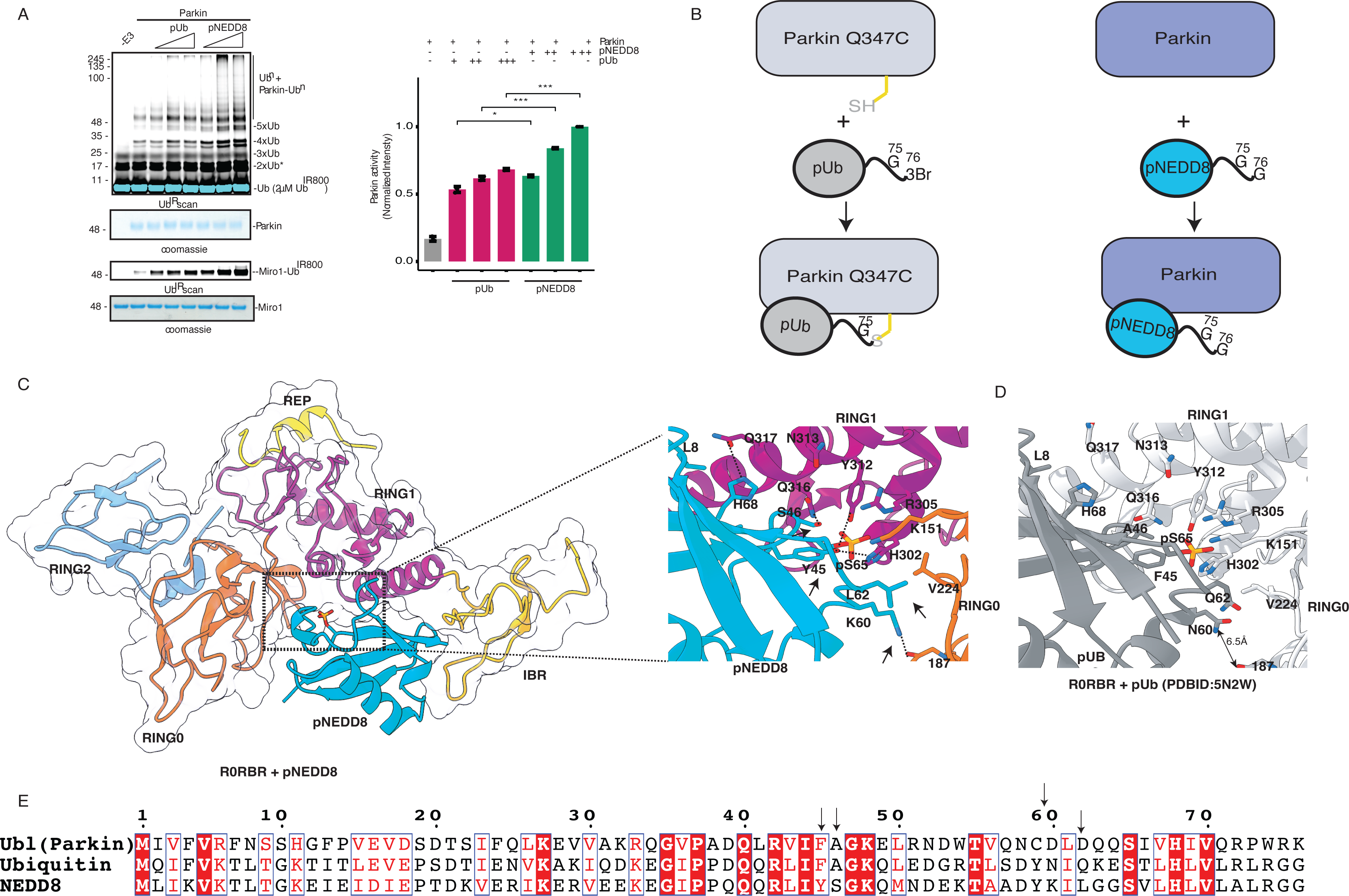
NEDD8 has evolved over ubiquitin to bind in the RING1 pocket and activate Parkin. A. Ubiquitination assay to test Parkin activation with increasing concentrations of pUb or pNEDD8. The lower panel shows Miro1 ubiquitination. Coomassie-stained gels showing Parkin/Miro1 were used as the loading controls. The ATP-independent band is indicated (*). The bar graph (left panel) shows the integrated intensities (mean ± s.e.m.) of ubiquitin chains from three independent experiments, as shown in the right panel. Statistical significance (\*\*\**P* < 0.001) was determined using a pairwise student’s t-test. B. Schematic representation showing the difference in approach to capture the crystal structure of Parkin and pUb ^22,26,30^ or pNEDD8 complex C. Key interactions between Parkin (colored domain-wise) and pNEDD8 (cyan). Unique hydrogen bonds between Parkin and pNEDD8 are highlighted with dashed lines. Favorable residues on NEDD8 that interact with the RING1 pocket are marked with arrows. D. Key interactions between Parkin (light gray) and pUb (gray). The distance between N60 of pUb and the carbonyl of the 187^th^ amino acid of Parkin is highlighted. E Sequence alignment of Ubl (Parkin), Ub, and NEDD8. Favorable residues on NEDD8 that interact with the RING1 pocket are marked with arrows.

Further, we wanted to understand the molecular details of the interactions between NEDD8 and Parkin. Unlike pUb bound human Parkin structures which were crystallized by introducing Q347C on Parkin and crosslinking with suicide probe of pUb (pUb-3Br) ^22,26,30^, we crystallized the complex of pNEDD8 with human Parkin without any modification of Parkin or pNEDD8 (Fig. 1B). The complex of human Parkin (R0RBR) and pNEDD8 was purified over size-exclusion chromatography and crystallized. The structure of Parkin and pNEDD8 complex was determined at 2.25 Å (Table 1).

The overall structure of the Parkin and pNEDD8 complex was quite similar to that of the Parkin and pUb complex (Extended Data Fig. 1A, 1B). Closer inspection of Parkin structures with pUb or pNEDD8 revealed some minor conformational changes in helix 1 (H1) on RING1 of Parkin. For example, Q317 is located away from the pUb interface in the pUb-bound Parkin structure, whereas Q317 is present at the interface of pNEDD8 and forms a hydrogen bond with H68 of pNEDD8 (Fig. 1C, 1D). In addition to the conserved interactions between Parkin and pUb or pNEDD8, new unique interactions are also observed between Parkin and pNEDD8. For example, S46 of pNEDD8 forms a hydrogen bond with Q316 of Parkin (Fig. 1C). Y45 of pNEDD8 forms hydrogen bonds with Y312 and H302 of Parkin (Fig. 1C). Furthermore, K60 of pNEDD8 forms a hydrogen bond with the carbonyl of L187 in Parkin (Fig. 1C). In addition to the above hydrogen bonds, L62 of pNEDD8 forms hydrophobic interactions with V224 of Parkin (Fig. 1C). Interestingly, among all known Ubls that can be phosphorylated by PINK1, the above residues are present only in pNEDD8 and are replaced by unfavorable residues in Ubl and Ub (Fig. 1E). Interestingly, assays performed with unphosphorylated NEDD8 or Ub also revealed higher activation of Parkin in presence of NEDD8 (Extended Data Fig. 1C). Overall, our data shows that NEDD8 is evolved over ubiquitin to bind more tightly in the RING1 pocket of Parkin.

### Parkin activators (pNEDD8/pUb) specifically bind to the RING1 pocket and cause allosteric conformational changes in RING2

Recent publications suggested that the RING0 pocket (comprising K211) binds to pUb leading to the feedforward loop of Parkin activation ^29,31^. These publications used constructs lacking RING2 and reported the binding of pUb on the RING0 and RING1 pockets. Our recent work demonstrated that RING2 negatively affects pUbl binding to RING0 ^30^. Therefore, we wondered whether the presence of RING2 would provide specificity for selecting Parkin activators in the RING0 pocket. To test this hypothesis, we used the untethered (TEV-treated) R0RBR (RING0-RING1-IBR-REP-RING2, the TEV site between IBR and REP) protein. We performed a RING2 displacement assay similar to our recent research, which showed RING2 displacement from untethered Parkin by pUbl/pParkin binding in trans ^30^. Notably, both pUb and pNEDD8 formed complexes with untethered R0RBR (Fig. 2A, Extended Data Fig. 2A). However, pUb/pNEDD8 binding failed to displace RING2 from RING0, as suggested by the coelution of REP-RING2 with R0RB (RING0-RING1-IBR) and pUb/pNEDD8 (Fig. 2A, Extended Data Fig. 2A), unlike pUbl/pParkin binding in trans, which displaces REP-RING2 from the Parkin core ^30^. Interestingly, the tighter binding of pNEDD8 (Fig. 1) also failed to displace REP-RING2 from the Parkin core (Extended Data Fig. 2A). Furthermore, the K211N mutation in the RING0 domain did not affect the complex formation between Parkin and pUb/pNEDD8 (Fig. 2B, Extended Data Fig. 2B). However, as expected, the H302A mutation in the RING1 domain disrupted the complex formation between Parkin and pUb/pNEDD8 (Fig. 2C, Extended Data Fig. 2C). Under none of the above conditions, unlike pUbl/pParkin binding in the RING0 pocket displacing the RING2 domain in trans ^30^, REP-RING2 was displaced from Parkin core, suggesting that pUb/pNEDD8 binds specifically to the RING1 pocket. These data also indicate that the movement of the RING2 domain after pUb binding is far less drastic than suggested earlier ^29,31^.

**Figure 2.**
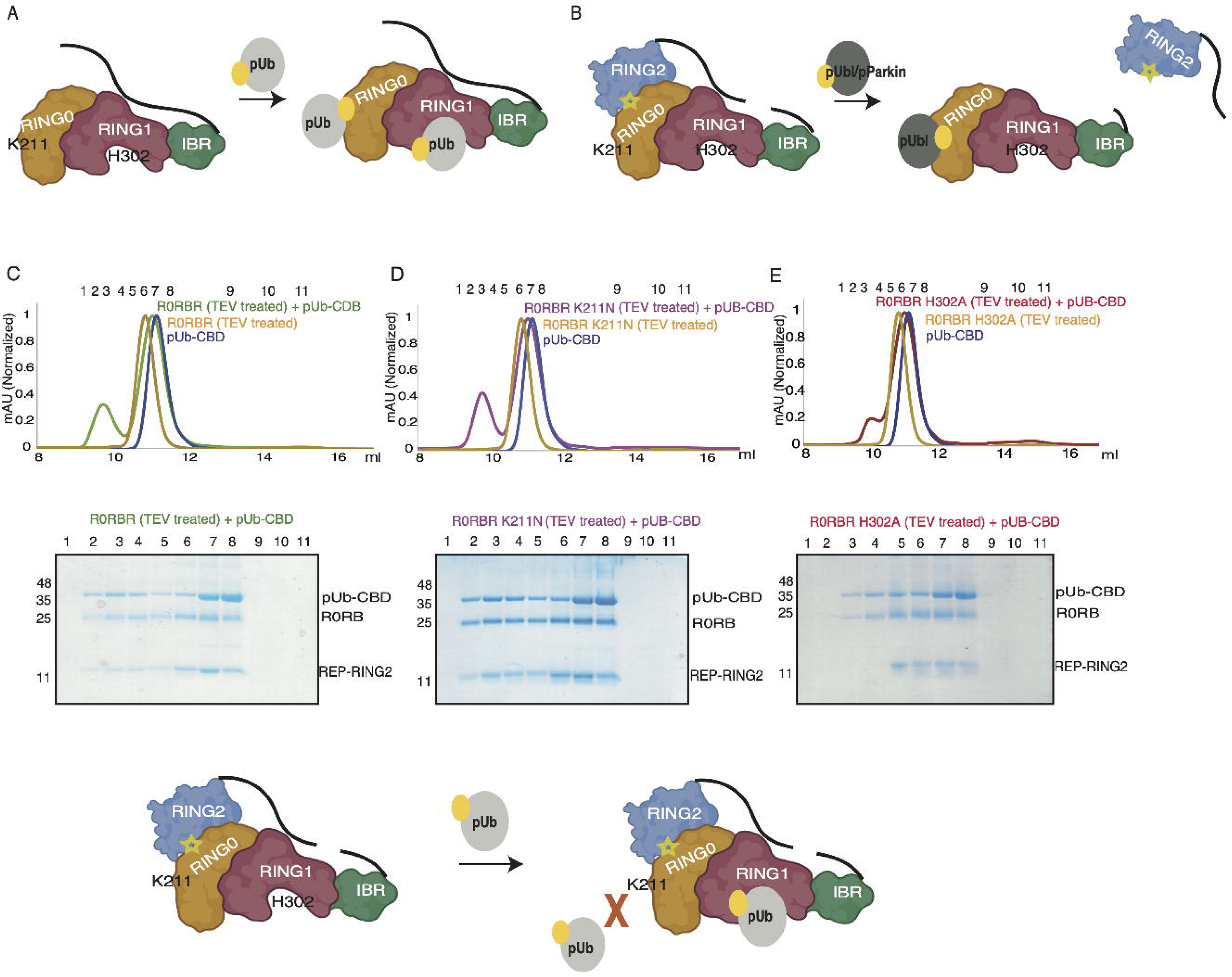
Size-exclusion chromatography (SEC) assay to check RING2 displacement and pUb binding. A. Schematic representation showing experimental design used previously ^29,31^ to capture pUb binding to RING0 and RING1 domains of Parkin B. Schematic representation showing experimental design used previously ^30^ to capture pUbl or pParkin binding to RING0 domain in trans. Catalytic C431 on RING2 is highlighted. C. SEC assay using Parkin (R0RBR, TEV-treated) and pUB (with a CBD tag). A colored key is provided for each trace. Coomassie-stained gels of the indicated fractions are shown. D. SEC assay using Parkin (R0RBR K211N, TEV-treated) and pUB (with a CBD tag). E. SEC assay using Parkin (R0RBR H302A, TEV-treated) and pUB (with a CBD tag). The lower panel is a schematic representation of the SEC analysis from panels A, B, and C.

Although RING2 was not completely displaced in the Parkin and pNEDD8 complex, we noticed significantly higher values of B-factor for RING2 and IBR domains in the pNEDD8 bound Parkin structure compared to the apo Parkin structure (Fig. 3). A higher B-factor indicates the flexibility of the RING2 region of Parkin due to pNEDD8 binding to the RING1 pocket. Flexible RING2 in the presence of pNEDD8 also explains the mechanism of Parkin activation due to allosteric conformational changes caused by phospho-NEDD8 binding in the RING1 pocket (Fig. 3). The flexibility in the RING2 leads to exposure of catalytic C431 on RING2 thus leading to activation of Parkin (Fig 3). Compared to the pNEDD8 bound Parkin structure, the pUb bound Parkin structure showed milder flexibility in the RING2 (Extended Data Fig. 3). Although lesser flexibility in the RING2 region of the pUb bound state of Parkin could be due to the cross-linking of pUb with the IBR domain of Parkin (Fig. 1B), weaker binding of pUb compared to pNEDD8, as suggested by crystal structure and biochemical data (Fig. 1), would also lead to a similar observation. These observations are consistent with the biophysical data in Fig. 2 & Extended Data Fig. 2, which did not show complete displacement of RING2 after binding pUb/pNEDD8. These data suggest the specific nature of the two pockets on Parkin and the unique conformational changes associated with these binding events. While pUb/pNEDD8 binding in the RING1 pocket leads to allosteric conformational changes resulting in flexible RING2, pUbl binding in RING0 leads to complete displacement of RING2 and higher activation of Parkin.

**Figure 3.**
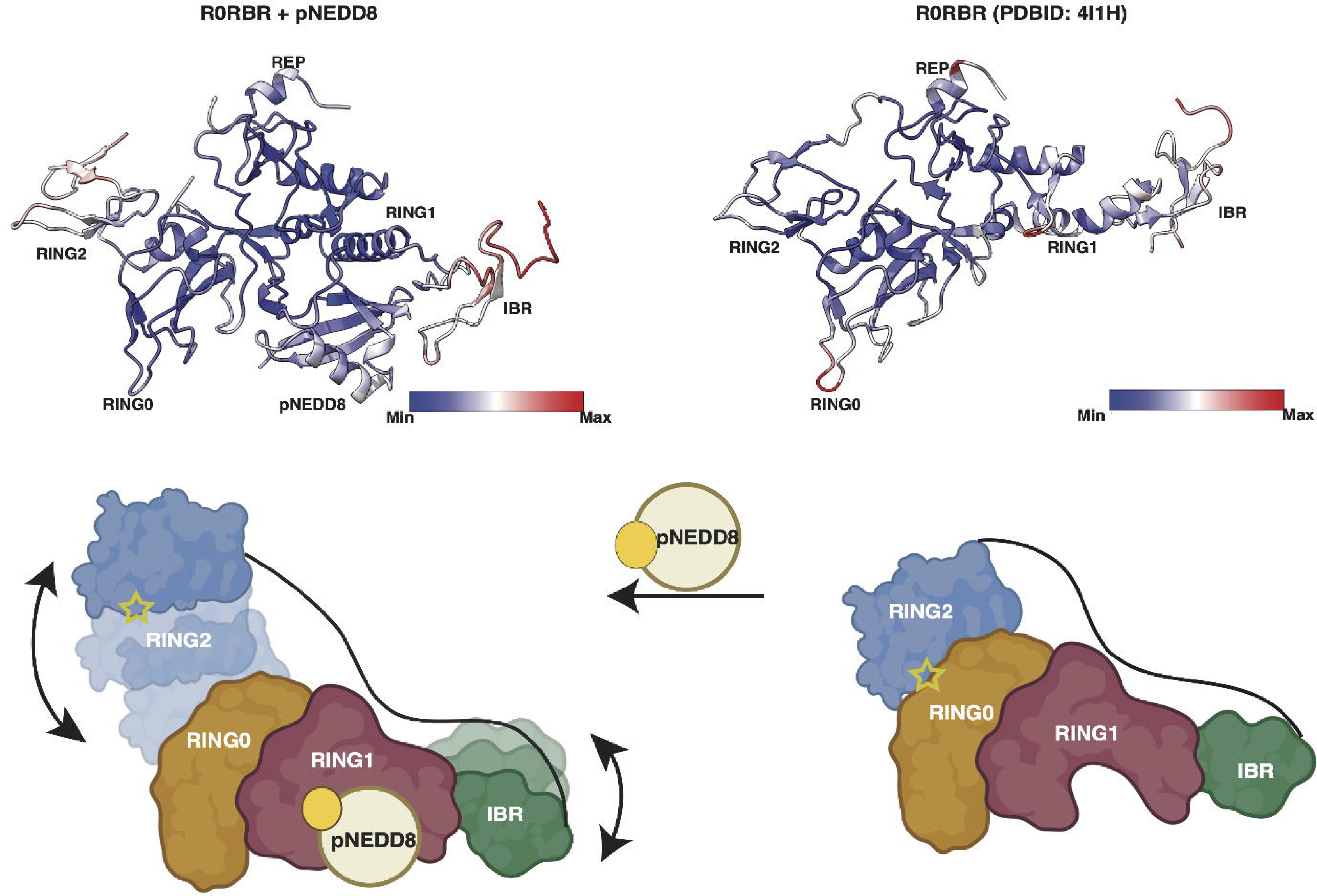
Crystal structure of Parkin and NEDD8 complex reveals flexible RING2. Residue-wise B-factor analysis of the Parkin (R0RBR) and pNEDD8 complex (left panel). For comparison, the right panel shows the residue-wise B-factor analysis of apo Parkin (PDBID: 4I1H). A scale is included to show the colorwise distribution of the B-factor, min (blue) to max (red). In the lower panel, the schematic representation highlights conformational changes (as per B-factor analysis) on Parkin due to the binding of pNEDD8. Catalytic C431 on RING2 is highlighted.

### Mechanism of specific interaction between Parkin and pUbl or pUb/pNEDD8

In contrast to the findings from previous studies ^29,31^, our data suggest that the two pockets (RING0 and RING1) on Parkin specifically recognize pUbl and pUb/pNEDD8. Although the sequence comparison in Fig. 1E suggested that key residues are conserved between Ubl, Ub, and NEDD8, differences at some key positions may help determine specificity. For example, L8 in Ub/NEDD8 is N8 in Ubl, which may not allow the binding of pUbl to the RING1 pocket (Fig. 1E & ^29^). Additionally, the difference between the critical residues mentioned in Fig. 1C-E helps determine the specificity of the RING1 pocket between pUb and pNEDD8. However, despite a very high sequence and structure similarity between pUbl, pUb, and pNEDD8, the inability of pUb/pNEDD8 to bind to the RING0 pocket (Fig. 2 & Extended Data Fig. 2) remains elusive.

Electrostatic surface charge analysis of Parkin, pUbl, pUb, and pNEDD8 revealed a distinct charge distribution on ubiquitin-like proteins/domain and Parkin (Fig. 4A-D). Surrounding of S65 of pUbl (Fig. 4A) are more negatively charged than pUb (Fig. 4B) or pNEDD8 (Fig. 4C). Furthermore, the regions surrounding the basic patch (K161, R163, K211) on the RING0 pocket were composed of electropositive groups (R104/R455) (Fig. 4D). Whereas, a negatively charged residue (E300) is positioned above the basic patch (K151, H302, R305) on the RING1 pocket of Parkin (Fig. 4D). The above positions face 60^th^ residue from pUbl and pUb or pNEDD8 bound to RING0 and RING1, respectively. Interestingly, D60 on Ubl was substituted with N60 or K60 on Ub or NEDD8, respectively (Fig. 1E & Fig. 4A-C). An electronegative group at the 60th position of Ubl would increase its interaction in the RING0 pocket (Fig 4A, 4D). Similarly, N60/K60 on Ub/NEDD8 would not favor interaction with the positively charged residues (R104/R455) around the basic patch of the RING0 pocket (Fig. 4B-D). We tested Parkin activation using pUb or pUb N60D to validate our structural analysis. Interestingly, compared to pUb, pUb N60D activated Parkin more robustly (Fig. 4E). The latter also suggests that pUb N60D binds to the RING0 pocket, which leads to a more significant displacement of RING2, causing higher activation of Parkin.

**Figure 4.**
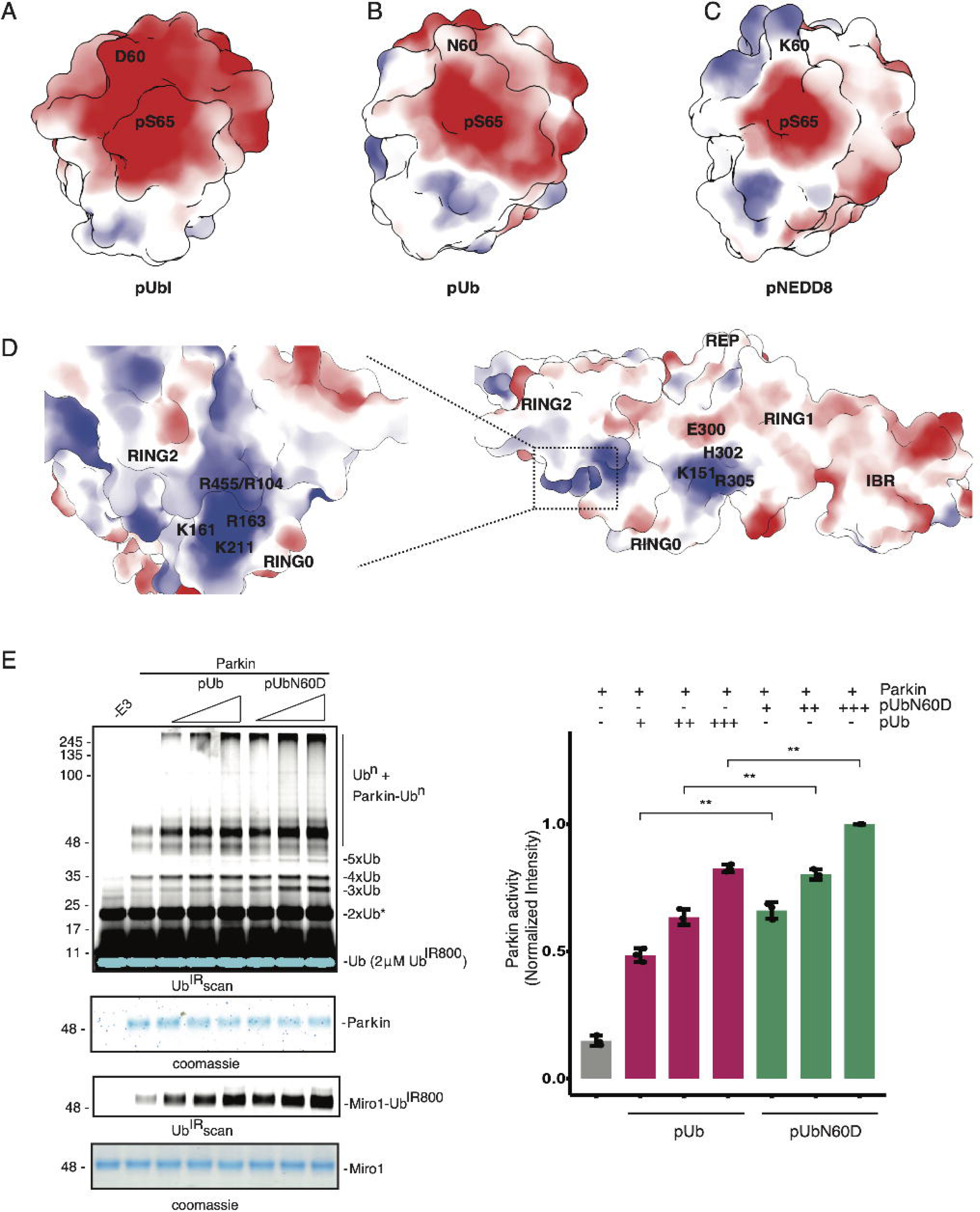
Unique charge distribution determines activators’ specificity on Parkin. A. The electrostatic surface charge distribution of the phospho-Ubl (pUbl) domain of Parkin. B. Electrostatic surface charge distribution of phospho-ubiquitin (pUb). C. The electrostatic surface charge distribution of the phospho-NEDD8 (pNEDD8) D. Electrostatic surface charge distribution around the basic patch on the RING0 and RING1 domains of Parkin. E. Ubiquitination assay to check Parkin activation with increasing concentrations of pUb or pUb N60D. The lower panel shows Miro1 ubiquitination. Coomassie-stained gels showing Parkin/Miro1 were used as the loading controls. The ATP-independent band is indicated (*). The bar graph (right panel) shows the integrated intensities (mean ± s.e.m.) of the ubiquitin chains from three independent experiments. Statistical significance (\*\*\**P* < 0.001) was determined using a pairwise student’s t-test.

To confirm the binding of pUb N60D to the RING0 pocket, we performed SEC assay using untethered Parkin. While native pUb binding did not lead to REP-RING2 displacement and resulted in binding only in the RING1 pocket (Fig. 5A), pUb N60D was able to bind on both pockets of Parkin, thus leading to a larger shift in the SEC assay (Fig. 5A). Furthermore, binding of pUb N60D with RING0 led to REP-RING2 displacement from Parkin core (Fig. 5A). To validate the latter, we tested binding of pUb N60D with K211N Parkin. K211N mutation led to the loss of pUb N60D binding to the RING0 pocket and thus did not displace REP-RING2 (Fig. 5B). However, H302A mutation perturbed the binding of pUb N60D to the RING1 pocket while binding to the RING0 pocket was intact which resulted in the displacement of REP-RING2 (Fig. 5B). Overall, this data establishes the mechanism of the specificity on the RING0 and RING1 pockets of Parkin.

**Figure 5.**
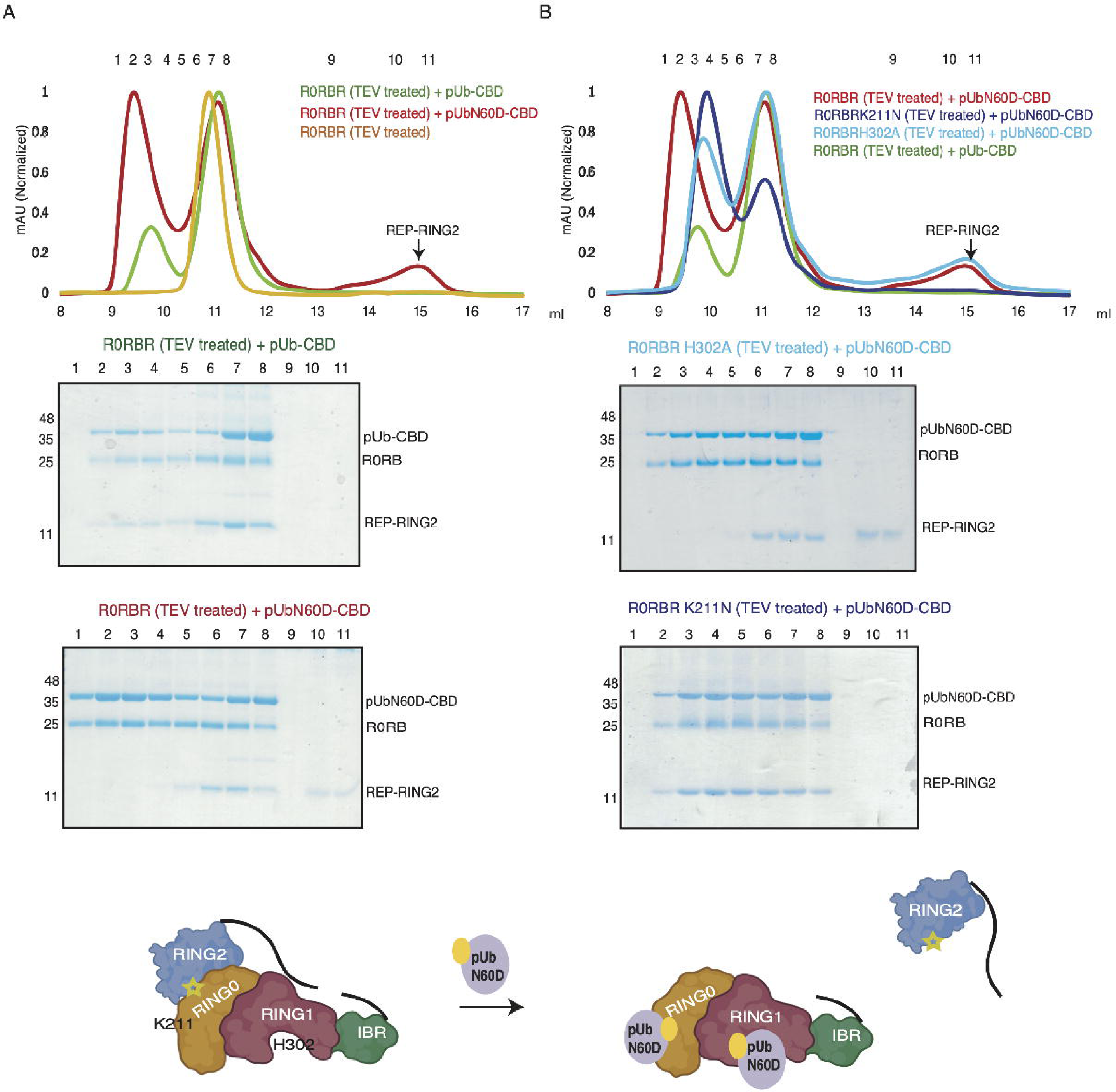
Size-exclusion chromatography (SEC) assay to check RING2 displacement and pUb N60D binding. A. SEC assay using Parkin (R0RBR, TEV-treated) and pUB/pUb N60D (with a CBD tag). A colored key is provided for each trace. The displaced REP-RING2 peak after binding of pUb N60D in the RING0 pocket of Parkin is marked with an arrow. Coomassie-stained gels of the indicated fractions are shown in the lower panel. B. SEC assay using Parkin (R0RBR, treated with TEV)/Parkin (R0RBR K211N, treated with TEV)/Parkin (R0RBR H302A, treated with TEV) and pUb/pUb N60D (with CBD-tag). A colored key is provided for each trace. The lower panel is a schematic representation of the SEC data analysis from panels A and B.

### Disease mutation K211N perturbs Parkin activity independent of pUb/pNEDD8 binding

The two basic patches on Parkin comprising H302 or K211 are suitable for binding with pSer65 of phosphorylated ubiquitin/NEDD8 or pUbl domain (Fig. 6A). Parkinson’s disease mutation K211N mutation on Parkin leads to loss of pUb-mediated Parkin activation suggesting interaction between K211 and pSer65 of pUb ^23,31^. However, our data suggest that the two pockets (comprising H302 or K211) of Parkin are specific, and pUb/pNEDD8 does not bind with the K211 pocket on RING0. We wanted to understand the role of the K211N mutation in Parkin activation. Additionally, as pNEDD8 activates Parkin more robustly than pUb (Fig. 1A), we wondered whether pNEDD8 would rescue the defect due to K211N mutation. Interestingly, although, unlike Parkin H302A, Parkin K211N does not exhibit any defect in its interaction with pNEDD8 (Extended Data Fig. 2), both Parkin K211N and Parkin H302A are 2-fold less active than native Parkin in the presence of pNEDD8 (Fig. 6B). This observation is consistent with the pUb-mediated Parkin activation on Parkin mutants (^31^ and Extended Data Fig. 4). We noticed that in the control lane without pNEDD8/pUb, the basal ubiquitination activity of Parkin K211N was lower than that of WT Parkin (Fig. 6B). Therefore, we decided to perform time course ubiquitination assay on WT/K211N Parkin in absence of pUb/pNEDD8. Interestingly, the K211N mutation led to an approximately 2-fold reduction in the basal level of Parkin activity that was independent of any activator (Fig. 6C). Overall, these data suggest that the loss of Parkin activity is not due to the inability of Parkin K211N to bind with pUb/pNEDD8 but rather due to some unknown reason.

**Figure 6.**
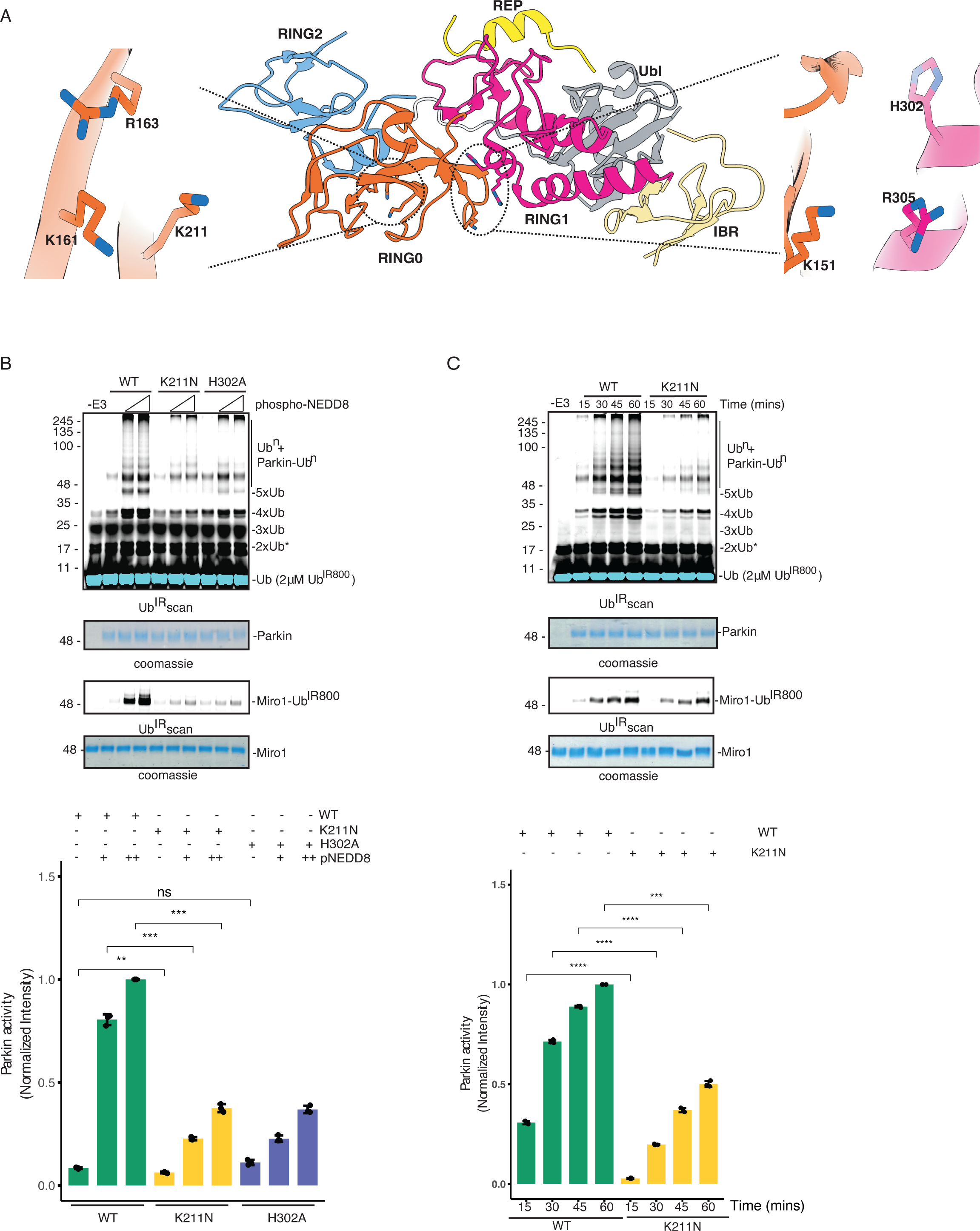
K211N mutation leads to loss of Parkin activity independent of pUb binding. A. Crystal structure of human Parkin (PDBID: 5C1Z) showing two pSer65 binding pockets on RING0 and RING1 domains. B. Ubiquitination assay to check WT/K211N/H302A Parkin activation with increasing concentrations of pNEDD8. The lower panel shows Miro1 ubiquitination. Coomassie-stained gels showing Parkin/Miro1 were used as the loading controls. The ATP-independent band is indicated (*). The bar graph (left panel) shows the integrated intensities (mean ± s.e.m.) of ubiquitin chains from three independent experiments, as shown in the right panel. Statistical significance (\*\*\**P* < 0.001) was determined using a pairwise student’s t-test. C. Time-course ubiquitination assay to compare the basal activity of WT Parkin and K211N Parkin in the absence of any activator (pUb/pNEDD8).

### Crystal structure of Parkin K211N and pNEDD8 complex reveals allosteric conformational changes that cause Parkin inhibition

As our data clearly showed that the loss of Parkin K211N activity was not due to the inability of Parkin to bind to pUb/pNEDD8 (Fig. 2, 6), we wanted to understand the molecular mechanism of the loss of Parkin K211N activity. To understand this, we crystallized and determined the structure of the R0RBR (K211N) and pNEDD8 complex. The structure of R0RBR (K211N) and pNEDD8 complex was determined at 1.8 Å (Table 1). Similar to R0RBR and pNEDD8 complex, structure of R0RBR (K211N) and pNEDD8 complex shows only one molecule of pNEDD8 bound to the RING1 pocket, and the overall Parkin conformation remained similar (Extended Data Fig. 5, Fig. 7A). Interestingly, unlike the flexible conformation of RING2 & IBR in the structure of native Parkin with pNEDD8 (Fig. 3), the lower B-factor of RING2 & IBR in the Parkin K211N structure confirmed that RING2 flexibility was reduced upon K211N mutation (Fig. 7A). Furthermore, closer inspection of structures revealed that due to K211N mutation, the RING0 conformation was changed and pushed toward RING2 (Fig. 7B, Extended Data Fig. 5). In mutant Parkin, N211 forms a hydrogen bond with R163 and pulls R163 away from K161 (Fig. 7B). The new conformation of R163 in Parkin K211N forces F209 away from its earlier position (Fig. 7B). In the new conformation, R163 forms hydrogen bonds with Q165, the carbonyl of D219, and N211 (Fig. 7B). Due to the above conformational changes, RING0 was pushed closer to the RING2 domain (Fig. 7B). These changes led to the formation of new hydrogen bonds between K435 (RING2) and the carbonyl of T177 (RING0), and between side chains of N428 (RING2) and S181 (RING0) (Fig. 7B). Additionally, in the Parkin K211N structure, K161 (RING0) forms a salt bridge with D460 (RING2). Furthermore, H433 (RING2) forms a hydrogen bond with W462 (RING2) in the Parkin K211N structure (Fig. 7B). The above interactions were absent in the WT Parkin structures, leading to an unlocked state of WT Parkin unlike the locked state seen in Parkin K211N (Fig. 7B). The new interactions between RING0 and RING2 in Parkin K211N would not allow displacement of RING2 and the exposure of C431, which is required for Parkin activation. Overall, our data shows that the K211N mutation leads to allosteric conformational changes in Parkin, restricting RING2 displacement and thus resulting in a loss of Parkin activity independent of pUb/pNEDD8 binding.

**Figure 7.**
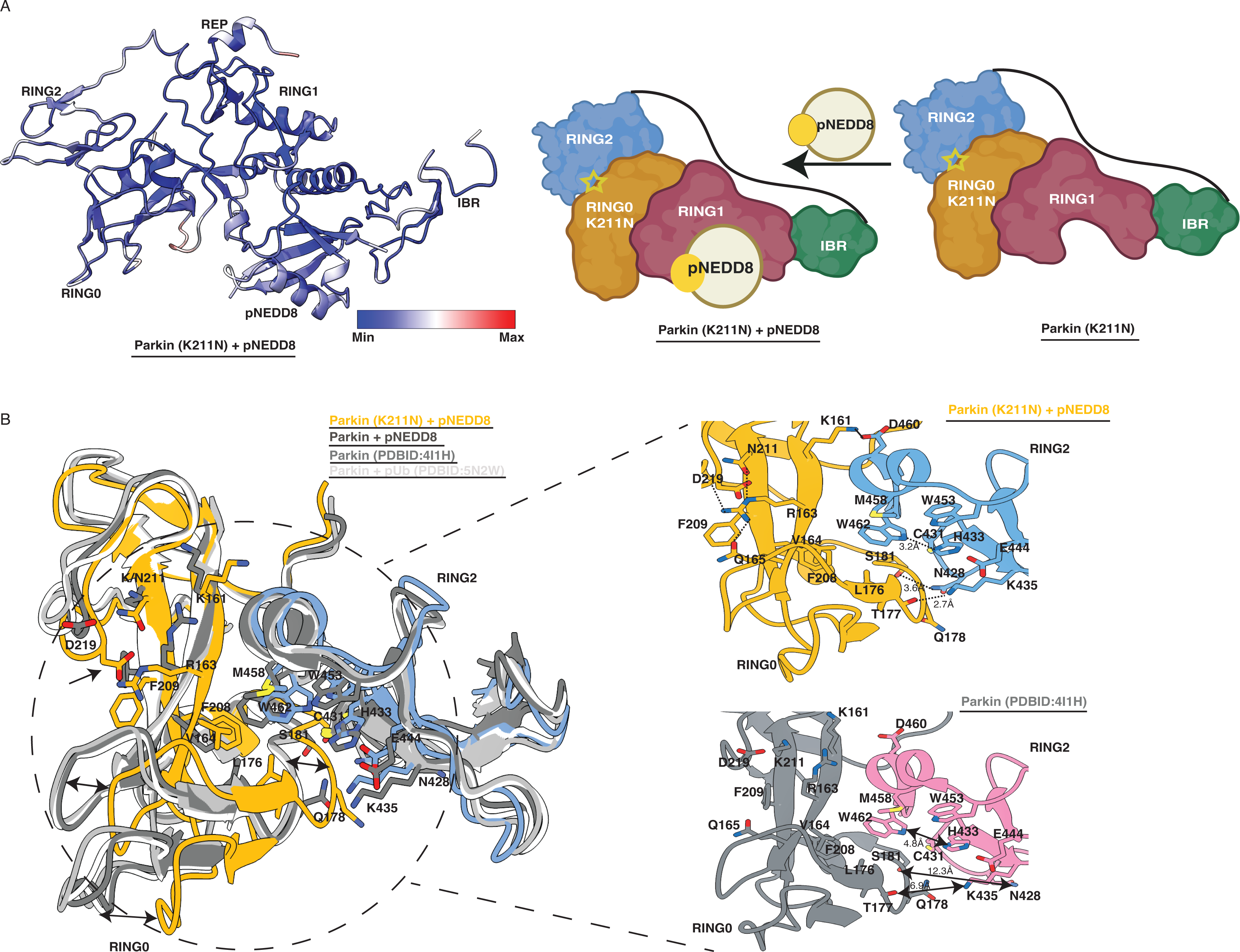
Crystal structure of Parkin (R0RBR K211N) and pNEDD8 complex reveals the mechanism of Parkin K211N inhibition. A. Residue-wise B-factor analysis of the crystal structure of Parkin (R0RBR, K211N) and pNEDD8 complex. A scale is included to show the colorwise distribution of the B factor, min (blue) to max (red). In the right panel, a schematic representation is used to highlight the conformation of Parkin (R0RBR, K211N) and pNEDD8 complex that is different from Parkin (R0RBR) and pNEDD8 complex (Fig. 3). Catalytic C431 on RING2 is highlighted. B. Overlay of native Parkin (apo or bound with pUb/pNEDD8) structures on K211N Parkin bound with pNEDD8. The RING0 and RING2 domains from various native Parkin structures are shown in shades of gray. The RING0 (orange) and RING2 (blue) domains of Parkin K211N are shown. Conformational shifts in the RING0 domain of Parkin K211N are highlighted with two-sided black arrows. The RING0 pocket with repositioned side chains (stick) due to K211N is highlighted with an arrow. Standalone structures of K211N Parkin and native Parkin are shown (right panel). Hydrogen bonds are shown as dashed lines, and the distance between the same atoms is highlighted with an arrow. The salt bridge between K161 (RING0) and D460 (RING2) is highlighted.

## Discussion

Mutations in the gene expressing the Parkin E3 ligase lead to a familial form of Parkinson’s disease (PD). Due to its key role, Parkin is a significant therapeutic target. In the last few years, several groups have extensively contributed toward understanding the mechanism of Parkin activation ^8–32^. Many Parkin structures have helped understand the mechanism of Parkin activation ^10,14,15,22–27,29–32^. The disease mutation K211N (on RING0), which leads to the loss of pUb-mediated Parkin activation ^23,31^, was an interesting phenomenon from a therapeutic perspective. Recently, a group suggested that pUb binds to the RING0 pocket and proposed a new feedforward mechanism of Parkin activation independent of pUbl ^29,31^. However, disease mutations on Parkin leading to loss of binding with its activators make therapeutic efforts of designing small-molecule Parkin activators even more challenging. Additionally, a lack of activators’ specificity may lead to unwanted catastrophic events in cells.

Unlike previous reports, we demonstrate that the RING0 and RING1 pockets specifically bind to pUbl and pUb/pNEDD8, respectively. pUb and pNEDD8 can bind to RING1; however, pNEDD8 shows robust binding and activation of Parkin than pUb. Structure of Parkin and pNEDD8 complex further confirms that pNEDD8 is preferred over pUb to bind in the RING1 pocket (Fig. 1 & 2). In contrast to previous reports ^29,31^, pUb/pNEDD8 do not bind to the RING0 pocket (Fig. 2 & Extended Data Fig. 2). The RING0 pocket of Parkin is recognized explicitly by the pUbl domain of Parkin, which could be utilized to activate Parkin in trans, as suggested by our recent work ^30^. Our data also shows how a unique charge distribution on pUbl/pUb/pNEDD8 and the two binding pockets of Parkin determine the specific recognition of Parkin activators (Fig. 4 & 5). Furthermore, the higher affinity of the RING1 pocket of Parkin for native NEDD8 than for native Ub (Extended Data Fig. 1C) explains the molecular mechanism of a previous finding, which showed that overexpression of the NEDD8 activation enzyme rescues the defects caused by silencing of PINK1 ^33^.

Conformational changes leading to Parkin activation remain elusive. Despite the more robust binding and higher activation by pNEDD8 than pUb (Fig. 1), RING2 was not completely displaced in the structure of pNEDD8 bound to Parkin (Fig. 3). Higher B-factor of RING2 in the crystal structure of Parkin and pNEDD8 complex suggests milder and allosteric displacement of RING2 after binding of activator in the RING1 pocket, unlike previous model suggesting complete RING2 displacement during pUb mediated Parkin activation ^29,31^. The crystal structure is also confirmed by biophysical assay, which shows that RING2 remained associated with the protein core after pUb/pNEDD8 binding (Fig. 2 Extended Data Fig. 2), unlike pUbl-mediated interactions with RING0 leading to complete displacement of RING2 ^30^. Furthermore, N60D mutation in ubiquitin, which mimics the Ubl domain of Parkin, results in binding between pUb N60D and the RING0 pocket, resulting in a complete displacement of RING2 from the core of Parkin and thus higher activation (Fig. 4 & 5). This study establishes that pUbl and pUb/pNEDD8, due to their specific binding to the two distinct pockets on Parkin, lead to larger/faster and smaller/slower displacements of RING2, respectively. The latter is also supported by the extent of Parkin activation due to different phosphorylation events.

K211N mutation in the RING0 pocket of Parkin was believed to perturb Parkin and pUb interaction ^29,31^. Herein, we demonstrate that the K211N mutation does not affect Parkin and pUb/pNEDD8 interaction (Fig. 2). Furthermore, we demonstrate that the K211N mutation leads to a loss of Parkin activity independent of pUb/pNEDD8 binding (Fig. 6). The latter is further confirmed by the Parkin K211N structure (Fig. 7). The structure of Parkin K211N shows allosteric conformational changes in the RING0 domain, which restricts the flexibility of RING2 by enhanced interactions between RING0 and RING2 (Fig. 7), thus leading to a loss of activity that is independent of pUb/pNEDD8 binding. The latter also explains the findings from a previous study, which showed that the K211N mutation reduces the recruitment of Parkin (which lacks the Ubl domain or Ser65) to mitochondria ^31^. However, unlike previously believed, the defect in Parkin recruitment was not due to the loss of interaction between pUb and Parkin K211N. Instead, the conformational changes that lead to a decrease in Parkin activity reduce ubiquitin chains and, thus, pUb levels, leading to less Parkin recruitment to mitochondria that is mediated by interactions between RING1 and pUb (Fig. 8). This is also supported by our recent study wherein under Parkin activity independent conditions H302A mutation completely abolished Parkin recruitment to the mitochondria ^30^. Meanwhile, under the same condition, K211N mutation mildly reduces Parkin recruitment, which is due to the interaction between Parkin molecules mediated by pUbl and RING0 in trans ^30^. This discovery would be helpful in the development of small-molecule activators against Parkin by pharmaceutical companies. Additionally, this study highlights the importance of performing a detailed structural analysis of various disease-related mutations on Parkin to aid in drug design against Parkinson’s disease.

**Figure 8.**
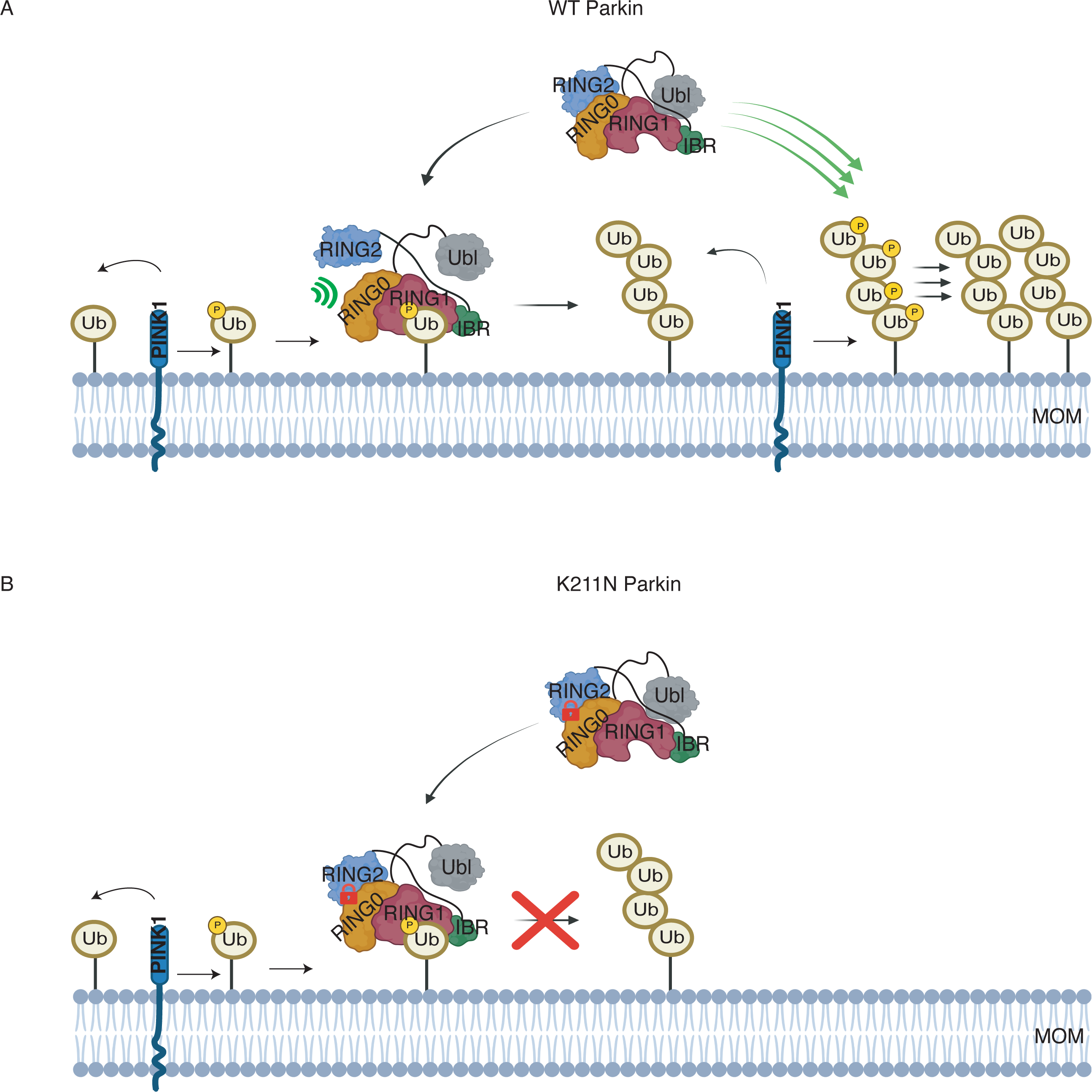
Model depicting the mechanism of pUb-mediated Parkin activation independent of pUbl phosphorylation. A. Mechanism of pUb binding mediated Parkin recruitment to mitochondria followed by feedforward activation is shown B. Mechanism of pUb binding mediated Parkin K211N recruitment to mitochondria and loss of feedforward loop is shown.

## Authors’ contributions

DRL performed all the experiments, solved the structures, and analyzed the data. SC purified various proteins. AK designed and supervised the research, refined the structures, analyzed the data, and arranged the funding. AK wrote the manuscript with input from DRL.

## Acknowledgments

We thank Prof. Helen Walden and Prof. Miratul M. Muqit for their useful discussions. We also thank Prof. Deepak Nair (RCB, Faridabad) and the Department of Biotechnology (Govt of India) for access to beamtime at the ESRF. We thank the ESRF (ID30B), Grenoble, France, and support staff for the data collection. The authors also acknowledge Shakti Dahe and other members of the AK group for helping with the various reagents. DRL is a PMRF fellow. AK is a recipient of the Innovative Young Biotechnology Award (DBT/12/IYBAl2019/03) and Ramalingaswami Fellowship (DBT/RLF/Re-entry/42/2019), which funded this project. AK also acknowledges IISER, Bhopal, and SERB (SERB/F/6520/2019-2020) for funding.

## Conflict of interest

The authors declare no conflicts of interest.

**Extended Data Figure 1. Analysis of the structure of Parkin and pNEDD8 complex**

A. Superimposition of the pNEDD8 (cyan) bound Parkin (R0RBR, different domains are colored) structure, and the pUB (gray) bound Parkin (gray) structure (PDBID:5N2W).

B. 2Fo-Fc map (gray) of the crystal structure of R0RBR (colored as panel A) and pNEDD8 (cyan) complex. The 2Fo-Fc map is contoured at 1.2 σ.

C. Ubiquitination assay to check Parkin activation with increasing concentrations of unphosphorylated Ub or NEDD8. The lower panel shows Miro1 ubiquitination. Coomassie-stained gels showing Parkin/Miro1 were used as the loading controls. The ATP-independent band is indicated (*). The bar graph (right panel) shows the integrated intensities (mean ± s.e.m.) of the ubiquitin chains from three independent experiments. Statistical significance (\*\*\**P* < 0.001) was determined using a pairwise student’s t-test.

**Extended Data Figure 2. SEC assay to check RING2 displacement and pNEDD8 binding**

A. SEC assay using Parkin (R0RBR, treated with TEV) and pNEDD8 (with a His-SUMO tag). A colored key is provided for each trace. Coomassie-stained gels of the indicated fractions are shown in the lower panel.

B. SEC assay using Parkin (R0RBR K211N, TEV-treated) and pNEDD8 (with a His-SUMO tag).

C. SEC assay using Parkin (R0RBR H302A, TEV-treated) and pNEDD8 (with a His-SUMO tag).

**Extended Data Figure 3. Residue-wise B-factor analysis of Parkin and pUb complex**

A scale is included to show the colorwise distribution of the B-factor, min (blue) to max (red).

**Extended Data Figure 4. Ubiquitination assay to check pUb-mediated activation of Parkin mutants**

Ubiquitination assays to check WT/K211N/H302A Parkin activation with increasing concentrations of pUb. The lower panel shows Miro1 ubiquitination. A Coomassie-stained gel showing Parkin/Miro1 was used as the loading control. The ATP-independent band is indicated (*).

**Extended Data Figure 5. Analysis of the structure of Parkin K211N and pNEDD8 complex**

Superimposition of various Parkin (both apo and in complex with pUb/pNEDD8) structures (shown in different shades of gray) with the pNEDD8 (green) bound R0RBR K211N (domains colored as shown) structure.

## Methods

### Molecular cloning

The human Parkin construct and several other constructs used in this study were made as per the details provided in our recent manuscript ^30^. Various Parkin mutations were generated using site-directed mutagenesis (SDM). A TEV protease site was introduced between the 382^nd^ - 388^th^ as explained in our recent manuscript ^30^. The NEDD8 gene was PCR amplified from cDNA from HEK393T cells and cloned in a pET15b vector with a His-SUMO tag. The Ub-Intein-CBD (Chitin-Binding Domain) construct was made by cloning the ubiquitin gene (residues 1-75) in the pTXB-1 vector with a His-tag (introduced after CBD using SDM). Pediculus humanus corporis PINK1 (115-575) was a gift from David Komander ^34^(Addgene plasmid # 110750). Ube1 was a gift from Cynthia Wolberger ^35^(Addgene plasmid # 34965).

### Protein expression and purification

Parkin constructs were expressed in *E. coli* BL21(DE3)pLysS cells. For Parkin (141-465) expression, the cells were grown until the OD reached 0.5-0.6, the temperature was decreased to 18°C, and the cells were induced with 50 µM IPTG supplemented with 400 µM ZnCl_2_. The cells were pelleted and lysed by sonication in lysis buffer (25 mM Tris pH 7.5, 300 mM NaCl, 5 mM imidazole, and 100 µM AEBSF). The lysate was clarified using centrifugation at 35,000 × g for 1 hour. His-tagged proteins were trapped using Ni-NTA resin. Unbound proteins were washed using wash buffer (25 mM Tris pH 7.5, 350 mM NaCl, and 10 mM imidazole). His-tagged proteins were eluted using elution buffer (25 mM Tris pH 7.5, 300 mM NaCl, 250 mM imidazole). His-SUMO was removed using the SENP1 protease. All the Parkin constructs were further purified on an Anion Exchange column (Hi-trap Q HP; GE Healthcare) followed by a Hiload 16/600 Superdex 200 pg column (GE Healthcare) preequilibrated with storage buffer (25 mM Tris pH 7.5, 75 mM NaCl, 0.25 mM TCEP). The SUMO-NEDD8 and ubiquitin-CBD constructs were expressed as described above without ZnCl_2_. After elution from Ni-NTA, the proteins were purified with a Hiload 16/600 Superdex 200 pg column (GE Healthcare) preequilibrated with storage buffer (25 mM Tris pH 7.5, 75 mM NaCl, 0.25 mM TCEP). The GST-PhPINK1, NEDD8, ubiquitin, Miro1, UbcH7, Ube1, and several other constructs used in this study were purified as described above and as mentioned in our recent manuscript ^30^.

### Ubiquitination Assay

Ubiquitination assays were carried out using full-length Parkin constructs. Reactions were performed at 25°C for 25 minutes in a buffer containing 25 mM Tris (pH 7.5), 50 mM NaCl, 10 mM MgCl2, 0.1 mM DTT, and 5 mM ATP. All the reactions contained 25 nM Ube1, 250 nM UbcH7 (E2), 1 µM E3, and 2 µM UbIR800. Ubiquitination of Miro1 was similar as above with 5 µM Miro1 and 0.5 µM Parkin at 25°C for 15 minutes. The ubiquitin labeling with IR800 dye was performed using sulfo-Cyanine7.5 maleimide (Lumiprobe Life science solutions) as per the manufacturer’s protocol ^36^, and UbIR800 was purified as described in our recent work ^30^. In Fig. 1A, Extended Data Fig. 1C, and Fig. 5E, reactions were carried out with increasing concentrations (1 µM, 2 µM, and 4 µM) of pNEDD8/pUb/pUbN60D/Ub/NEDD8. Fig. 4B and Extended Data Fig. 4 show the results obtained with 1 µM and 2 µM pNEDD8/pUb. Reactions were stopped by adding SDS dye and heating at 95°C for 5 mins. The samples were loaded on a gradient gel and resolved using 1X MOPS running buffer. The gels were scanned using a Li-COR® Odyssey Infrared Imaging System. All the reactions were repeated at least 3 times and quantified using ImageJ software ^37^. The first auto-ubiquitination event on Parkin was used for quantification, as shown in various figures. The statistical analysis was performed using R.

### Purification of phosphorylated proteins

GST-PhPINK1 was used for the phosphorylation of ubiquitin, ubiquitin (N60D), and NEDD8 in buffer containing 50 mM Tris (pH 8.5), 100 mM NaCl, 10 mM MgCl2, 10 mM DTT, and 10 mM ATP buffer at 25°C for four hours. Completion of the phosphorylation reaction was verified using PhoS-Tag-Acrylamide (FUJIFILM). PhPINK1 was trapped using GSH resin, and the flowthrough containing phosphorylated proteins was purified over Hiload 16/600 Superdex 75 pg column (GE Healthcare) preequilibrated with buffer containing 25 mM Tris (pH 7.5), 75 mM NaCl, and 0.25 mM TCEP.

### Protein crystallization

Parkin R0RBR (141-465) or R0RBR K211N (141-465) were purified similarly as above up to ion exchange chromatography using a Hitrap Q FF (GE Healthcare) column. Parkin was incubated with a 2-fold molar excess of phospho-NEDD8, and the complex was purified with a Hiload 16/600 Superdex 75 pg column (GE Healthcare) preequilibrated with buffer containing 25 mM Tris (pH 7.5), 75 mM NaCl, and 0.25 mM TCEP. Crystals of the R0RBR complex were obtained in 0.2 M sodium sulfate, 0.1 M Bis-Tris propane 7.5, and 20% w/v PEG 3350 at 4°C. Crystals of the R0RBR K211N complex were obtained in 0.1 M MMT 8.0 and 25% w/v PEG 1500 at 18°C. R0RBR and R0RBR K211N crystals with pNEDD8 were vitrified using their mother liquor containing 20% ethylene glycol and 20% PEG 400, respectively.

### Data collection and structure determination

The data were collected at ID30B of the European Synchrotron Radiation Facility (ESRF), Grenoble, France. The data were processed using XDS ^38^ and Xia2 ^39^ for R0RBR and R0RBR K211N, respectively. Aimless in the CCP4 suite ^40^ was used for data scaling. Structures were determined using Phaser ^41^. R0RBR (PDB id: 4I1H) and NEDD8 (PDB id: 1NDD) were used as search models. The models were built in Coot ^42^ and refined in Refmac5 ^43^. The data were deposited in the protein data bank, and the PDB codes are included in Extended Data Table 1.

### Size-exclusion chromatography assay

All the binding experiments were performed using a Superdex 75increase 10/300 GL column (GE Healthcare). Five hundred microliters of reaction mixture containing 20 µM phospho-Ub (CBD tag)/phospho-UbN60D (CBD tag)/phospho-NEDD8 (His-SUMO tag) was incubated with 10 µM R0RBR (treated with TEV and depleted of TEV)/R0RBR K211N (treated with TEV and depleted of TEV)/R0RBR H302A (treated with TEV and depleted of TEV) for 30 minutes on ice before injection. Fractions from each run were analyzed over SDSLPAGE.

