## Supplementary figures and images for "Specific interactions with phospho-Ubls and allosteric conformational changes regulate the E3 ligase activity of Parkin"

### Extended Data Figure 1

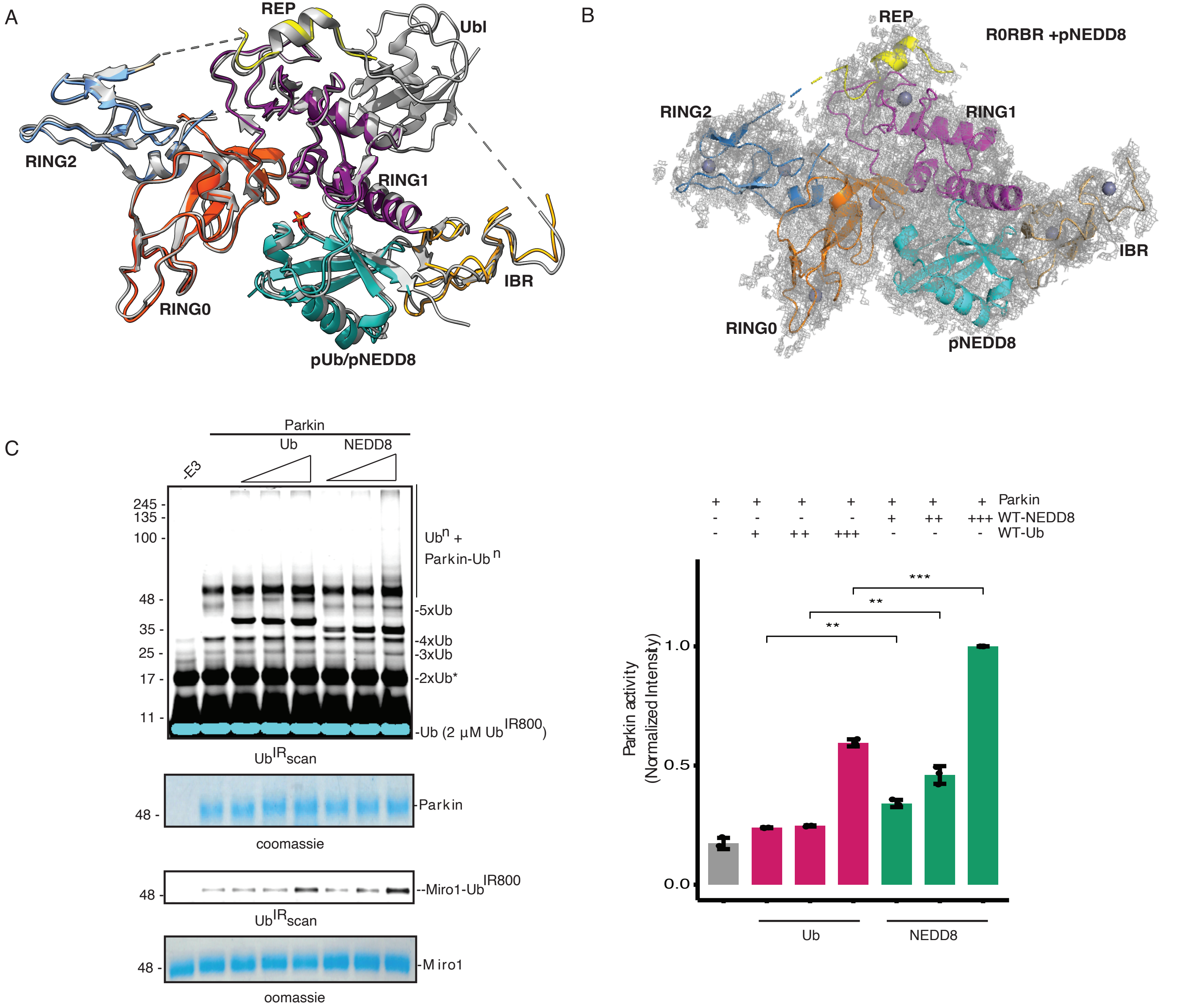


Extended Data Figure 1

### Extended Data Figure 2

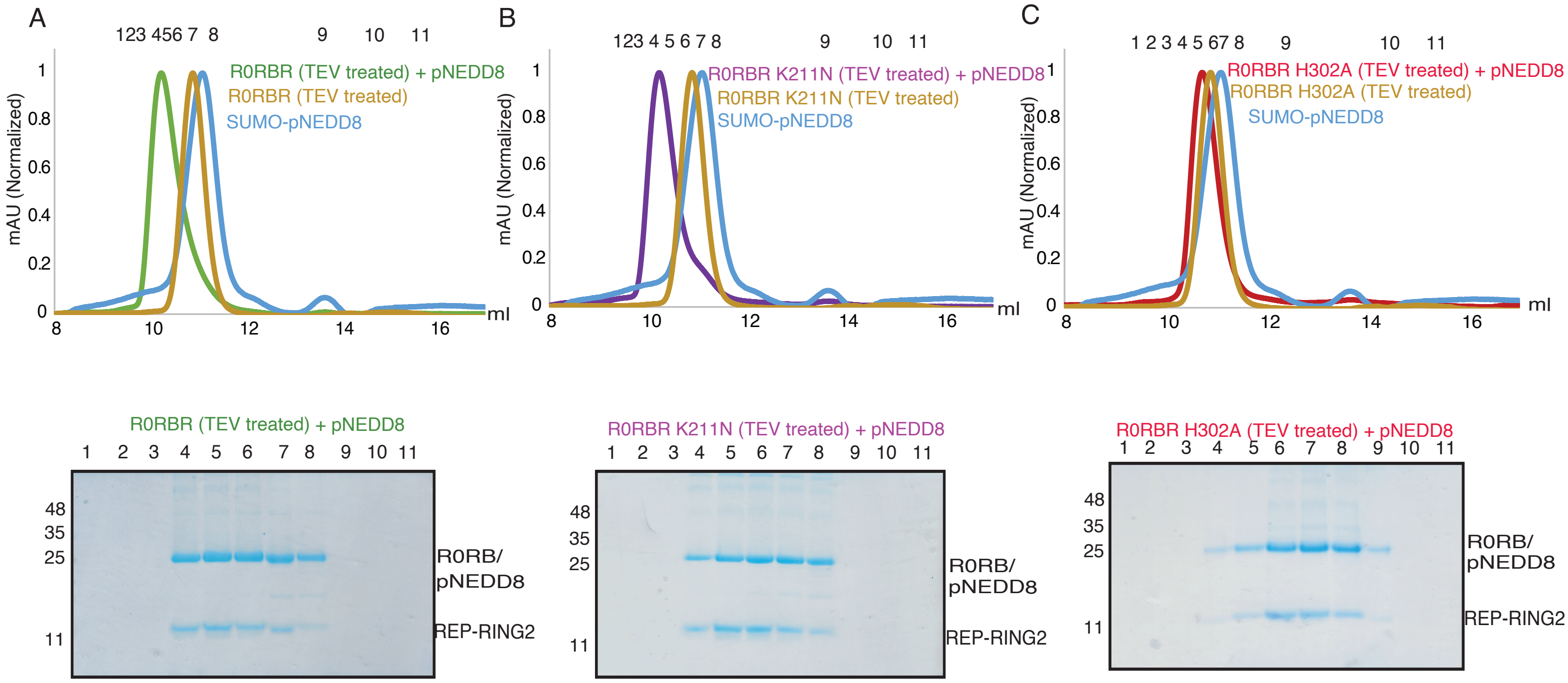


**Extended Data Figure 2**

### Extended Data Figure 3

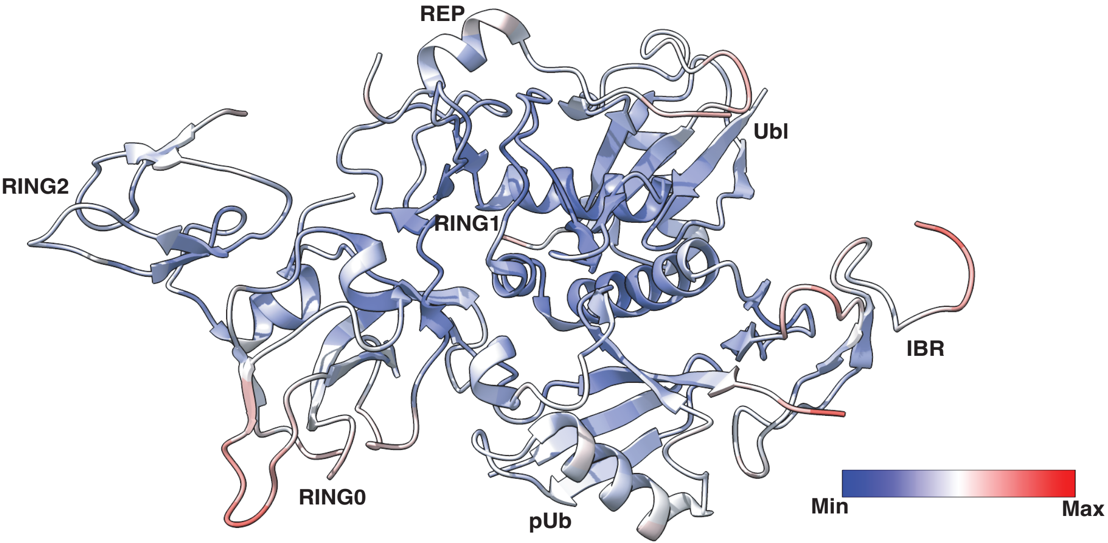


Parkin + pUb (PDBID: 5N2W)

Extended Data Figure 3

### Extended Data Figure 4

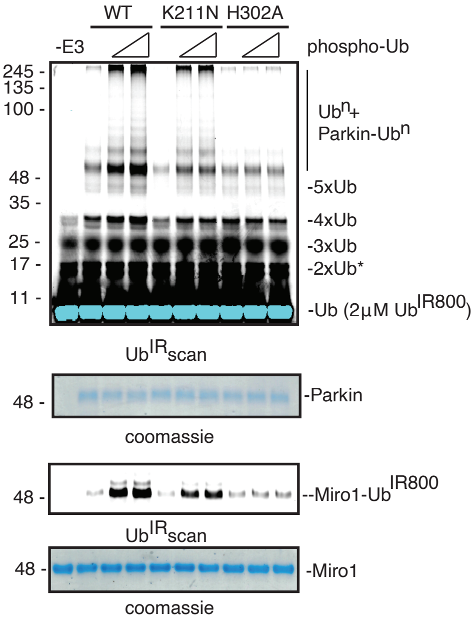


#### Extended Data Figure 4

### Extended Data Figure 5

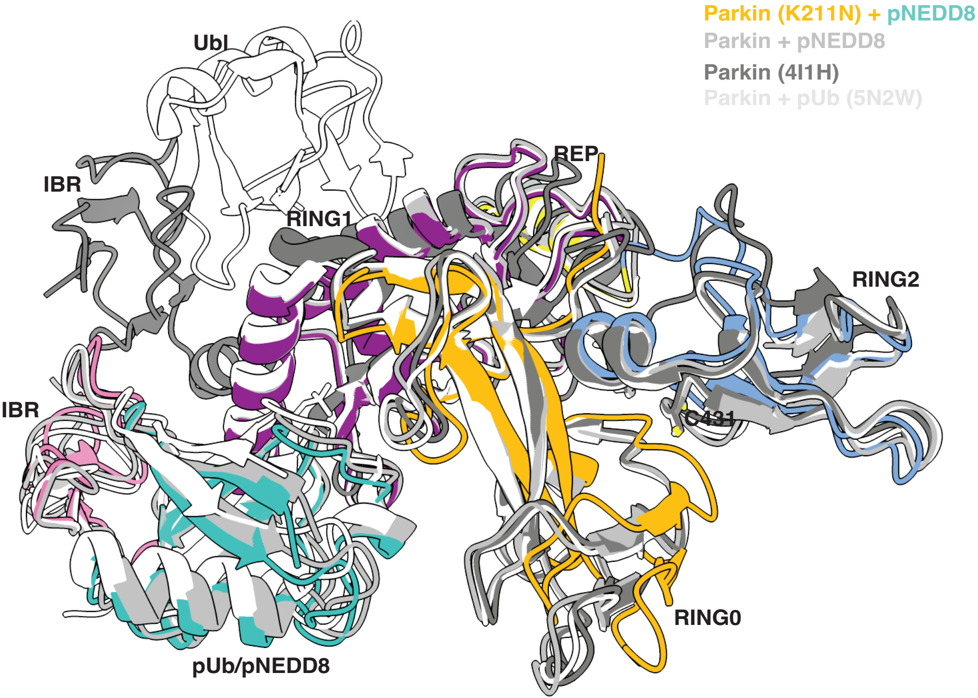


Extended Data Figure 5
